# Nucleosome fragility is associated with future transcriptional response to developmental cues and stress in *C. elegans*

**DOI:** 10.1101/047860

**Authors:** Tess E. Jeffers, Jason D. Lieb

## Abstract

Nucleosomes have structural and regulatory functions in all eukaryotic DNA-templated processes. The position of nucleosomes on DNA and the stability of the underlying histone-DNA interactions affect the access of regulatory proteins to DNA. Both stability and position are regulated through DNA sequence, histone post-translational modifications, histone variants, chromatin remodelers, and transcription factors. Here, we explored the functional implications of nucleosome properties on gene expression and development in *C. elegans* embryos. We performed a time-course of micrococcal nuclease (MNase) digestion, and measured the relative sensitivity or resistance of nucleosomes throughout the genome. Fragile nucleosomes were defined by nucleosomal DNA fragments recoverable preferentially in early MNase-digestion time points. We found fragile nucleosomes at locations where we expected to find destabilized nucleosomes, like transcription factor binding sites where nucleosomes compete with DNA-binding factors. Contrary to our expectation, the presence of fragile nucleosomes in gene promoters was anti-correlated with transcriptional activity. Instead, genes with fragile nucleosomes in their promoters tended to be expressed in a context-specific way, operating in neuronal response, the immune system, and stress response. Nucleosome fragility at these promoters was strongly and positively correlated with the AT content of the underlying DNA. There was not a strong correlation between promoter nucleosome fragility and the levels of histone modifications or histone variants. Our data suggest that in *C. elegans* promoters, nucleosome fragility is primarily a DNA-encoded feature that poises genes for future context-specific activation in response to environmental stress and developmental cues.

## INTRODUCTION

The fundamental unit of eukaryotic chromatin is the nucleosome, which consists of 147 bp of DNA wrapped around an octamer of histone proteins (Luger et al. 1997). Nucleosomes have important structural and regulatory functions in organizing the genome and restricting access of regulatory factors to the DNA sequence (Henikoff 2008). As such, the interactions between nucleosomes and DNA strongly influence the regulation of gene expression by determining DNA accessibility for transcription factors and RNA polymerase. In addition to regulated nucleosome assembly and disassembly through the action of histone chaperones and chromatin remodelers, nucleosome stability is influenced by histone modifications, histone variants, DNA features encoded in *cis*, and competition with DNA-binding factors in *trans*. For the purposes of this manuscript, we define “stability” qualitatively, as the propensity of a given nucleosome to remain intact and at that position, rather than to be evicted, disassembled, or translocated to a substantially different position. A complete picture of the mechanisms governing nucleosome stability is fundamental to understanding how gene expression is dynamically regulated.

Nucleosome stability has been studied *in vitro* using sensitivity to enzymatic digestion or salt concentration (Bloom and Anderson 1978; Simon and Felsenfeld 1979; Burton et al. 1978; Li et al. 1993; Jin and Felsenfeld 2007; Wu and Travers 2004; Polach and Widom 1995; 1999; Anderson et al. 2002). Genome-wide adaptations of these methods have been used to identify nucleosome position and stability *in vivo*. More recent studies in yeast, *Drosophila*, plants, and mammals have used varying concentrations of the enzyme micrococcal nuclease (MNase) to identify nucleosomes with differential sensitivity to MNase digestion *in vivo* (Xi et al. 2011; Henikoff et al. 2011; Chereji et al. 2015; Kent et al. 2011; Weiner et al. 2010; Vera et al. 2014; Lombraña et al. 2013; Kubik et al. 2015). Nucleosomes sensitive to low concentrations of MNase have been labeled as “fragile”, and have been associated with transcription factor binding sites (Vera et al. 2014), active origins of replication (Lombraña et al. 2013), gene promoters (Xi et al. 2011), and genomic sequences with high AT content (Chereji et al. 2015). Thus, both DNA-encoded sequence features and active processes in *trans* influence nucleosome fragility. The relationship between fragility and nucleosome function remains unclear. For example, one study reported fragile nucleosomes at the promoters of repressed stress-response genes during normal growth (Xi et al. 2011), while another found fragile nucleosomes at the promoters of highly transcribed genes in yeast (Kubik et al. 2015).

We performed a timecourse of MNase digestion in *C. elegans* mixed-stage embryos to study the relationship between nucleosome fragility and gene activity in a developing multicellular organism. In our study, fragile nucleosomes were associated with lowly expressed genes and genes expressed in a context-specific fashion. Although we found that competition with *trans* factors promoted nucleosome fragility, our data suggest that the majority of highly fragile nucleosomes in the *C. elegans* embryo are not due to *trans* factors but rather to *cis* features encoded in the DNA sequence. Together, our data indicates that the fragility of nucleosome-DNA interactions may aid in poising genes for induction in response to stress or developmental cues.

## RESULTS

### A digestion timecourse identifies nucleosomes with differential MNase sensitivity

We postulated that functionally distinct nucleosomes in the *C. elegans* could be distinguished by the length of time it took them to be liberated from bulk chromatin by MNase digestion. Previous studies using this approach defined nucleosomes released early in the timecourse as “fragile” and those released later in the timecourse as “resistant” (Xi et al. 2011). To identify nucleosomes of differential sensitivity genome-wide, we isolated mixed-stage embryos from *C. elegans*, treated them with formaldehyde to cross-link the chromatin, isolated nuclei, and digested the chromatin with MNase (**Figure 1A**). After 2, 4, 8, 15, and 30 minutes of digestion we removed a chromatin aliquot and performed paired-end Illumina sequencing on the mononucleosomal fragments liberated at each time point (**Figure 1B**). Without MNase, chromatin remained intact and undigested. After addition of the enzyme, a stereotypic chromatin ladder rapidly formed, and a small proportion of total chromatin became mononucleosomal. As digestion proceeded, the mononucleosomal fraction increased while polynucleosomal fractions were depleted (**Figure 1C**). We performed two replicate experiments on native chromatin and two replicate experiments on formaldehyde-fixed chromatin samples. Results from the native and fixed chromatin were very similar (**Supplemental Figure 1**). We therefore focused our downstream analysis on fixed chromatin for maximum compatibility with previously-generated datasets.

**Figure 1.**
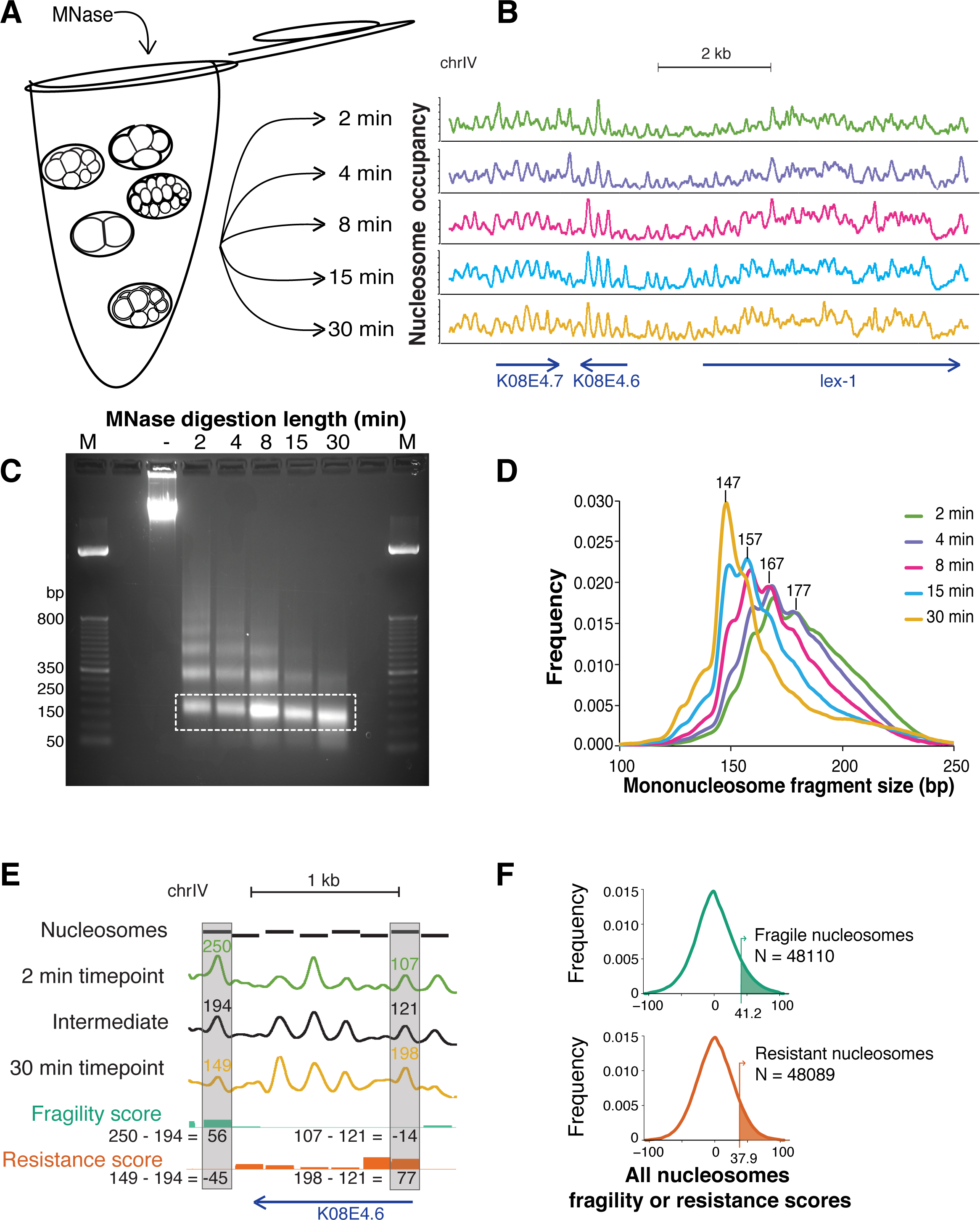
An MNase digestion time course on *C. elegans* embryos. **(A)** Mixed-stage embryos were collected from gravid hermaphrodites by bleach treatment. Dissociated nuclei from mixed-stage embryos were incubated with MNase for 2, 4, 8, 15, or 30 minutes. **(B)** Paired-end reads from each timepoint were mapped to the *C. elegans* genome, normalized, and Gaussian smoothed for display. High signals represent regions of the genome protected from MNase digestion. Region plotted: chr IV position 12,074,951 to 12,084,347. **(C)** Representative image of an N2 embryo MNase digestion timecourse after gel electrophoresis. For each timepoint, mononucleosome-sized fragments were excised from the gel (white box) and used for paired-end Illumina DNA sequencing. Size markers (M) are indicated. **(D)** Mononucleosome fragments are shorter with increasing MNase digestion time, in 10 bp increments. **(E)** Calculation of fragility and resistance scores. Fragility: for each nucleosome, the average occupancy of the intermediate timepoints is subtracted from the 2 minute timepoint. Resistance: for each nucleosome, the average occupancy of the intermediate timepoints is subtracted from the 30 minute timepoint. Intermediate timepoints are 4, 8, and 15 minutes. Region plotted: chr IV position 12,076,980 to 12,078,364. (**(F)** Distribution of fragility and resistance scores at all nucleosomes. The top 10% of each class (shaded in green and orange, respectively) were considered “Fragile” or “Resistant”.

Mononucleosomal DNA fragments released earliest during the digestion were larger (median size of 2-minute nucleosomal fragments: 180 bp) than fragments released later in the timecourse (median of 30-minute nucleosomes: 155 bp) (**Figure 1D**). Among the digestion timepoints, nucleosome sizes decreased in 10 bp increments, reflecting the MNase digestion preference for WW (AA, AT, TA, or TT) dinucleotides and the 10-11 bp periodicity of the DNA helical turn (McGhee and Felsenfeld 1983; Deniz et al. 2011; Trifonov and Sussman 1980; loshikhes et al. 2011) (**Figure 1D**). This is consistent with the model that with increasing lengths of digestion time, MNase will cleave long linkers and any unwrapped ends of nucleosomal DNA. Although the genome-wide occupancy profiles of mononucleosomal fragments were globally similar across the timepoints (**Figure 1B**, **Supplemental Figure 2**), there were a number of substantial differences in the nucleosome maps among the timepoints (**Supplemental Figure 3**, **Figure 1E**).

To systematically study nucleosomes of differential sensitivity to MNase, we assigned each nucleosome both a fragility and a resistance score (**Supplemental Figure 4**). For each timepoint, we first called nucleosome positions and then assigned each nucleosome an occupancy score (see Methods for details). The fragility score for a nucleosome is defined by subtracting the average occupancy score of the intermediate timepoints (4, 8, and 15 min) from the occupancy score of the 2 min timepoint. Conversely, a resistance score is computed by subtracting the average occupancy score of the intermediate timepoints from that of the 30 min timepoint (**Figure 1E**). Thus, fragility and resistance scores were generally reciprocal to each other at a given nucleosome, but not necessarily so. We defined the top 10% of nucleosomes with the highest fragility or resistance scores as “fragile” or “resistant” nucleosomes, respectively (**Figure 1F**).

### *Trans-factors* increase nucleosome fragility

We sought to address whether nucleosome fragility was a consequence of competition with DNA binding proteins and other *trans* factors. Trans-acting factors disrupt nucleosomes by competing with histones for binding to the DNA sequence (Simpson 1990; Adams and Workman 1995). We first examined regions of the genome where we expected to find nucleosomes destabilized by competition with other DNA-binding factors, for example at transcription factor binding sites (TFBS). We collected a set of 35,062 TFBS bound at any stage of *C. elegans* development, as identified by transcription factor (TF) ChIP-seq from the modENCODE consortium (Araya et al. 2014). TFBS in the *C. elegans* genome on average show strong affinity to histones *in vitro* (Locke et al. 2013). A nucleosome occupancy model based solely on DNA sequence also predicted *C. elegans* TFBS to be nucleosome bound (Kaplan et al. 2009) (**Figure 2A**). *In vivo*, however, these sites show a dip in nucleosome occupancy, consistent with the footprint of TF binding. Moreover, TFBS had high fragility scores on average (**Figure 2B**). These data are in agreement with previous reports from yeast to humans that transcription factors compete with nucleosomes for access to DNA (Barozzi et al. 2014; Wang et al. 2012; Ozonov and van Nimwegen 2013). To further investigate the relationship between TF binding and fragility we broke TFBS into groups depending on the number of TFs bound at a site. Although the majority of TFBS identified in *C. elegans* are bound by a single factor, some sites may be bound by many different TFs (Chen et al. 2014; Araya et al. 2014; Boyle et al. 2014). Fragility scores increased as a function of the number of TFs bound at a single TFBS (**Figure 2C**).

**Figure 2.**
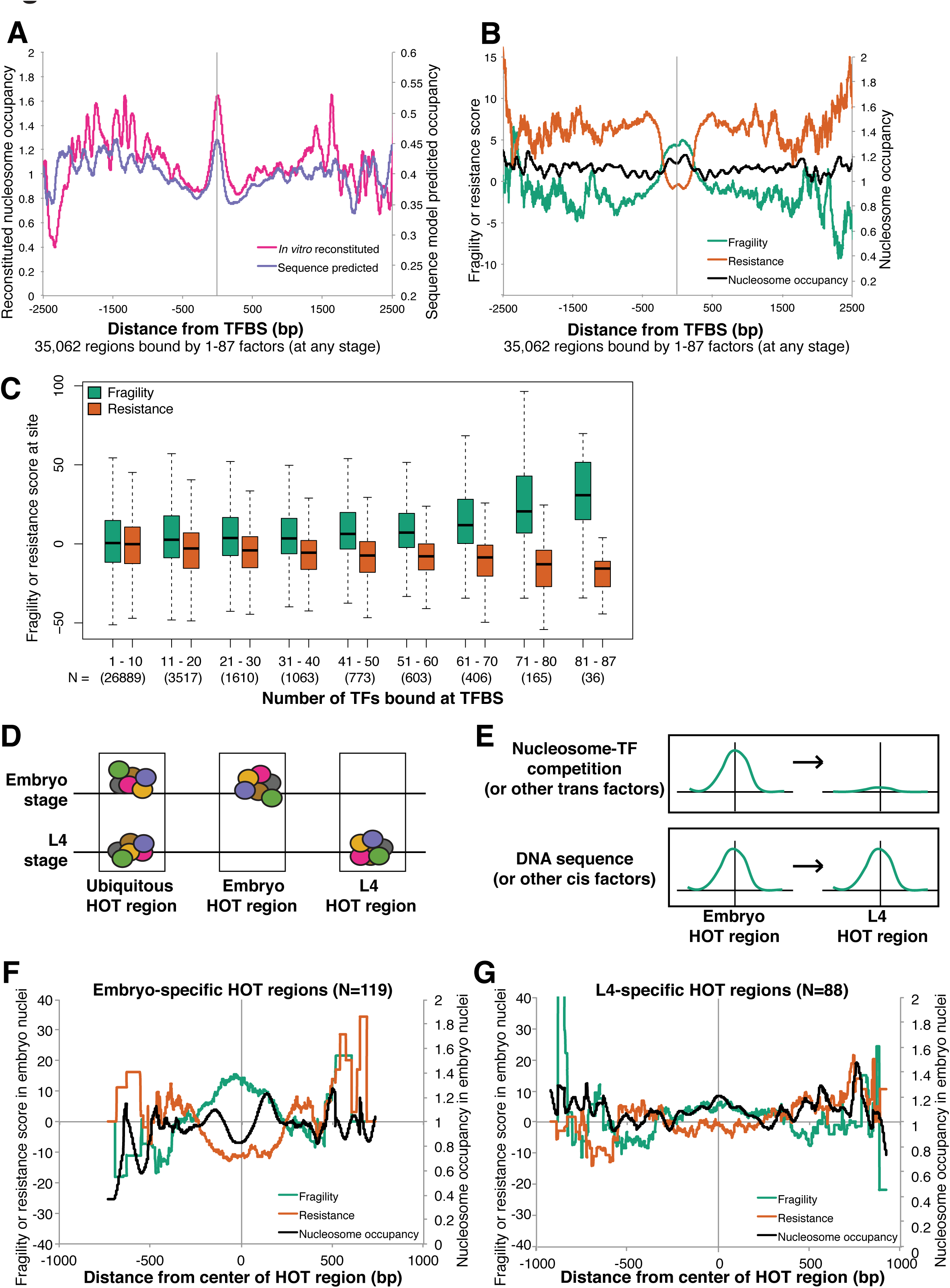
Competition with transcription factors influences nucleosome fragility. **(A)** Average reconstituted nucleosome occupancy scores (Locke et al. 2013) and computational nucleosome occupancy model scores (Kaplan et al. 2009) at 35,062 regions bound at any stage by any number of transcription factors. **(B)** Average fragility, resistance, and intermediate nucleosome occupancy scores are plotted around the same set of intervals from **(A)**. **(C)** Nucleosome fragility scores are higher at sites bound by more transcription factors. Boxplot of average fragility or resistance scores at groups of sites bound by different numbers of transcription factors. N = number of regions in each category. **(D)** Cartoon characterization of how embryo-specific and L4-specific HOT regions were identified. Embryo HOT region: binding sites bound only in the embryo. L4 HOT region: binding sites bound only in the L4 stage. **(E)** Model to distinguish whether *trans* (**top**) or *cis* (**bottom**) effects result in nucleosome fragility at a given nucleosome in the embryo. Hypothetical fragility scores are represented. (**(F)** Fragility, resistance, and nucleosome occupancy scores measured in the embryo at 119 regions found to be specifically HOT in the embryo. (G) Fragility, resistance, and intermediate nucleosome occupancy scores at 88 regions found to be specifically HOT in the L4 stage worm.

We found that TFBS had high fragility scores despite their intrinsic preference for nucleosome formation *in vitro*. One possible explanation is that transcription factors destabilize nucleosomes at their binding sites, causing the fragility at TFBS. Alternatively, TFBS may contain DNA sequences that disfavor nucleosome formation *in vivo*, thereby increasing nucleosome fragility. To distinguish among these possibilities, we identified a set of TFBS specifically bound at different developmental stages (**Figure 2D**). We hypothesized that if active competition with TFs increases nucleosome fragility, then TFBS bound by TFs only in the embryo should be fragile in embryos, whereas TFBS bound only in the L4 larval stage should not be fragile in embryos (**Figure 2E**, top). Alternatively, if DNA sequence influences nucleosome fragility then the embryo-specific and L4-specific TFBS should be equally fragile in embryos (**Figure 2E**, bottom). Due to their high fragility scores and dynamic nature, we focused our analysis on HOT regions, TFBS where significant enrichments (false discovery rate <5%) in multiple transcription factor binding sites are observed (Araya et al. 2014). We found that embryo-specific HOT regions had high nucleosome fragility and low nucleosome occupancy (**Figure 2F**, **Supplemental Figure 5**). In contrast, L4-specific HOT regions showed lower nucleosome fragility in embryonic samples and higher nucleosome occupancy than embryo-specific HOT regions (**Figure 2G**). These results support the hypothesis that active competition with transcription factors *in vivo* contributes to nucleosome fragility despite their intrinsically nucleosome favoring properties *in vitro*.

### Nucleosome fragility increases throughout heat-shock genes upon induction

The preceding analysis found a correlation between transcription factor binding and nucleosome fragility. We next sought to test the relationship between fragile nucleosomes and *trans* factors more explicitly. Moderate transcription levels have been shown to cause displacement and repositioning of nucleosomes in gene bodies (Reinberg and Sims 2006). At extremely highly transcribed genes, such as heat shock responsive genes after induction, it has been proposed that RNA Pol II molecules occupy the entire gene body (Merz et al. 2008; Cole et al. 2014; Schwabish and Struhl 2004; Kristjuhan and Svejstrup 2004). We hypothesized that nucleosome fragility should increase at gene bodies after inducing high levels of transcription, as a result of nucleosome competition with transcribing RNA polymerase II (Pol II). To test whether we could induce nucleosome fragility, we designed a heat shock experiment in conjunction with an MNase-seq timecourse (**Figure 3A**).

**Figure 3.**
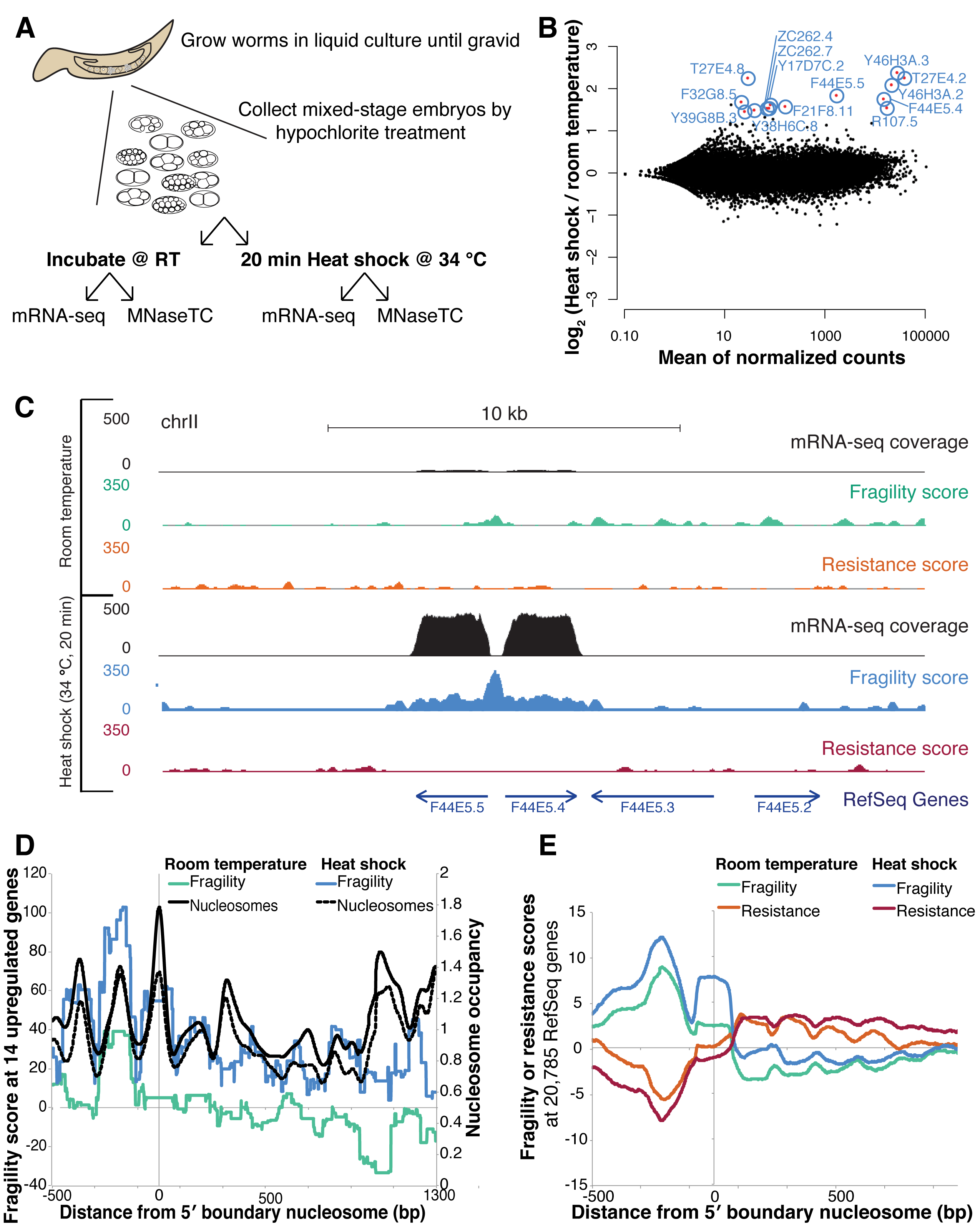
Heat shock increases nucleosome fragility at the promoter and gene body of upregulated genes. **(A)** Experimental overview. Mixed-stage embryos were either incubated at room temperature (RT) or heat shocked at 34°C (HS) for 20 minutes. Subsequently, embryos were fixed and used for an MNase-seq timecourse or stored in TRIzol and used for RNA-seq. **(B)** mRNA-seq of differentially expressed genes after a 20 minute HS at 34°C. Significantly differentially expressed genes (p_adj_ < 0.1) shown in red. **(C)** Increased fragility scores after heat shock in the coding region of F33E5.4 and F33E5.5, two divergently transcribed hsp-70 orthologues. Region plotted: chr II position 11,749,925 to 11,770,394. **(D)** Nucleosome fragility and nucleosome occupancy at 11 significantly differentially expressed genes with and without heat shock. **(E)** Nucleosome fragility and nucleosome resistance at all 20,768 coding genes with and without heat shock.

Heat shock in *C. elegans* activates HSF-1 and HSF-2, two homologues of the mammalian HSF1 transcription factor, which bind heat shock elements (HSE) in the promoters of heat shock-responsive genes to upregulate their expression (Åkerfelt et al. 2010). Upon heat shock, the heat-shock-response genes undergo rapid chromatin remodeling and colocalize with the nuclear pore complex (Rohner et al. 2013). Using RNA-seq, we identified 14 genes that are rapidly upregulated after a brief (20 minute) heat shock at 34 °C (**Figure 3B**, **Supplemental Figure 6**). We then analyzed how fragility scores changed at those genes after heat shock (**Supplemental Figure 7**). Though nucleosome occupancy remained largely unchanged, we found nucleosome fragility dramatically increased both 5′ and 3′ of heat-shock genes, as well as in the gene body itself (**Figure 3C, 3D**). Notably, promoter and +1 nucleosome fragility increased on average genome-wide, although gene-body fragility was specific to the set of heat shock-induced genes (**Figure 3E**). This result suggests that the rapid induction of these genes increases nucleosome competition with Pol II. This mode of transcription has been suggested to remove the entire histone octamer, rather than FACT-mediated H2A-H2B recycling (Kulaeva et al. 2010; Kireeva et al. 2002). Alternatively, previous studies of the Hsp70 locus in *Drosophila* have shown that heat shock induces rapid transcription-independent loss of gene-body nucleosomes (Petesch and Lis 2008). Future experiments with transcription inhibitors may clarify the exact mechanism by which gene body nucleosomes become fragile after heat shock. Our results demonstrate that nucleosome fragility can be modulated by *trans*-acting factors like transcription factors and RNA polymerase II, and is not solely dependent on DNA sequence.

### Nucleosome fragility near genes is anti-correlated with expression

We found high fragility scores at the types of genomic locations where destabilized nucleosomes had been previously reported, namely transcription factor binding sites and the gene bodies of newly induced genes (Urnov and Wolffe 2001). We investigated the genome-wide distribution of fragile nucleosomes (nucleosomes with the highest 10% of fragility scores; **Figure 1F**) in detail. Fragile nucleosomes were enriched 5′ and 3′ of genes, specifically at the promoter ‐2, ‐1, and +1 nucleosomes, and at the terminal nucleosome (TN) and TN+1 nucleosomes (**Figure 4**, **Supplemental Figures 1, 8**). Resistant nucleosomes (nucleosomes with the highest 10% of resistance scores (**Figure 1F**) were enriched in gene bodies (**Supplemental Figures 1, 8**).

**Figure 4.**
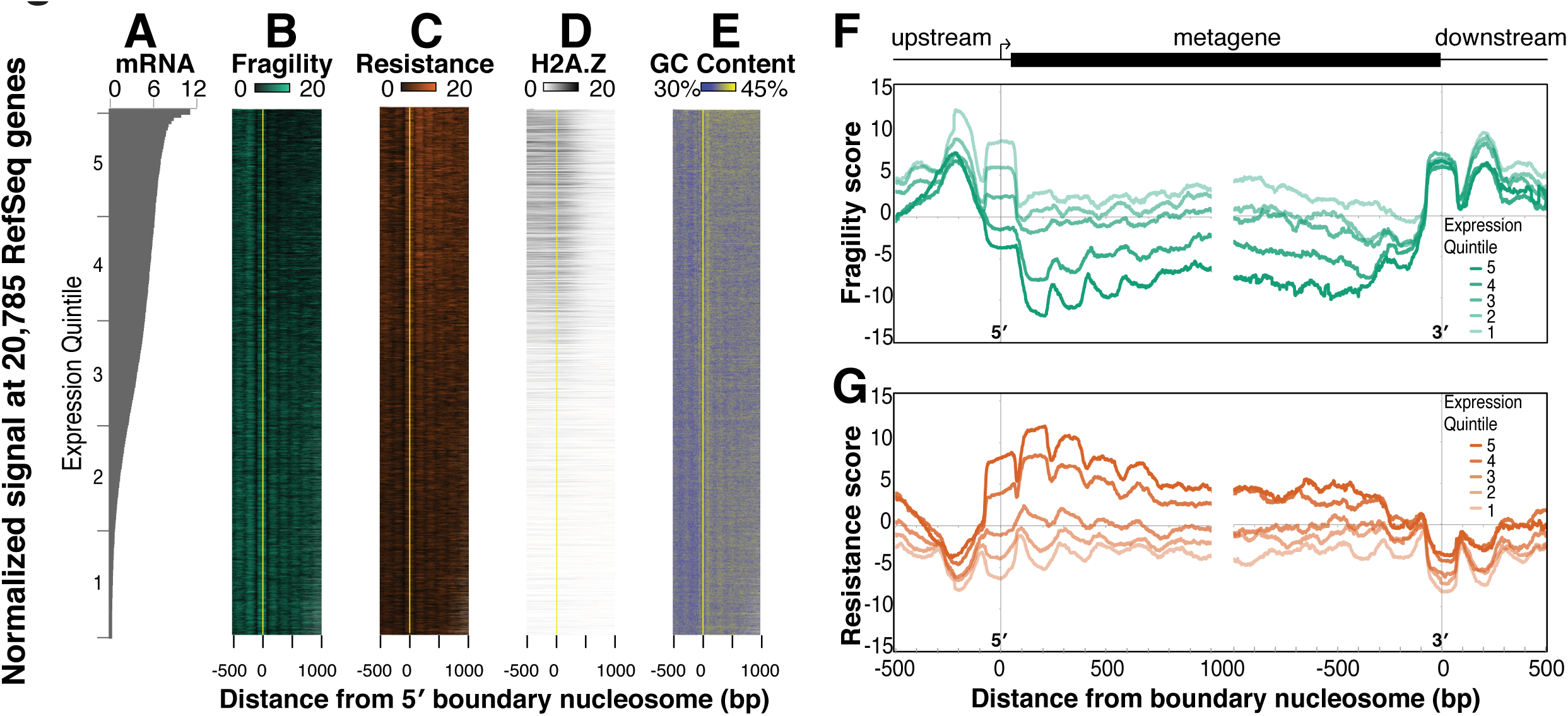
Fragility is enriched 5′ and 3′ of genes, and on average is anti-correlated with gene expression. **(A)** Log_2_ DESeq-normalized number of read counts measured by mRNA-seq at 20,785 genes, ordered by their relative expression. **(B)** Heatmap of fragility scores (green) at. Genes were aligned by the center of the first nucleosome downstream from the transcript start site, known as the +1 or 5′ Boundary Nucleosome (yellow line). **(C)** Same as in **(B)**, except resistance scores are plotted in red. **(D)** Same as **(B)**, except for HTZ-1 input-normalized ChIP-seq signals (Ho et al Nature 2014). **(E)** Same as **(B)**, except the average GC content (as a percentage of 100%) in 5 bp windows is plotted. (**(F)** Fragility and (G) resistance scores around the 5′ and 3′ boundary nucleosomes averaged over expression quintiles (highest expressed 20% in dark orange or green, lowest expressed 20% in lightest orange or green). Quintile 1: 0 to 4.5 normalized counts. Quintile 2: 4.5 to 65. Quintile 3: 65 to 619. Quintile 4: 619 to 2209. Quintile 5: > 2209.

With increasing distance from the promoter, nucleosomes become less sharply positioned (Mavrich et al. 2008; Yuan et al. 2005). To investigate whether the higher nucleosome resistance scores at highly expressed genes were a consequence of more consistent nucleosome positioning, we asked whether fragility or resistance scores were correlated with the standard deviation of the nucleosome center, or “fuzziness”. Neither nucleosome fragility nor nucleosome resistance scores were correlated with the fuzziness of the nucleosome at the intermediate timepoint (fragility vs. fuzziness: R = 0.03, resistance vs. fuzziness: R = ‐0.06), suggesting that the susceptibility or resistance to MNase digestion is not a direct consequence of more or less well-positioned nucleosomes (**Supplemental Figure 9**).

Given the positive relationship between fragility and transcription factor binding (**Figure 2C**), and given that induction resulted in increased nucleosome fragility throughout heat-shock genes (**Figure 3C, 3D)**, we expected that nucleosome fragility would be enriched in the promoters or gene bodies of highly expressed genes in the embryo, which exhibit high levels of TF and Pol II binding as measured by ChIP-seq (**Supplemental Figure 10**) (Ho et al. 2014). To our surprise, gene expression levels on average were anti-correlated with nucleosome fragility at both promoters and gene bodies (**Figures 4A, 4B, 4F**, **Supplemental Figure 12**. R = ‐0.17) and positively correlated with nucleosome resistance (**Figures 4A, 4B, 4G**, **Supplemental Figure 12**. R = 0.11). For example, in contrast to our earlier observation at the heat shock genes, we failed to find a correlation between ongoing expression and nucleosome fragility at the gene bodies of the most highly transcribed genes (compare **Figure 3D** to **Supplemental Figure 11**). Perhaps only newly-induced genes display gene-body fragility, or extremely high levels of transcription are required to induce fragility in gene bodies.

Although overall nucleosome fragility scores were high 5′ and 3′ of all genes, including at the majority of TFBS (**Figure 2**), fragile nucleosomes occurred preferentially at the promoters of lowly-expressed genes (**Figures 4B, 4F**, **Supplemental Figure 10**). No single transcription factor profiled in the embryo significantly overlapped the distribution of fragile nucleosomes (**Figure 1F**, **Supplemental Figure 10**). Though previous reports have suggested that the histone variant H2A.Z may act to promote nucleosome instability (Jin and Felsenfeld 2007; Jin et al. 2009; Xi et al. 2011), we did not observe a significant overlap between previously-identified H2A.Z-containing nucleosomes (Ho et al. 2014) and fragile nucleosomes (**Figures 4D**, **Supplemental Figure 13**). These data collectively suggest that two separate mechanisms account for nucleosome fragility depending on the genomic context. In places where nucleosomes are directly in competition with transcription factors (**Figure 2**) or in the bodies of exceptionally highly expressed or newly induced genes (**Figure 3**) fragility arises through competition with transcription factors or other DNA-associated proteins. By contrast, fragility 5′ and 3′ of genes at locations with few TF binding events appears to be determined by another mechanism, which we explored next.

### Nucleosome fragility is correlated to cis-encoded DNA features

We hypothesized that *cis* features may be responsible for the fragility of nucleosomes at the promoters of lowly-expressed genes. We examined the DNA sequences occupied by fragile and resistant nucleosomes, and compared these to sequences occupied by all nucleosomes in the genome (**Figure 5**). Compared to the set of all nucleosomes, DNA sequences occupied by fragile nucleosomes had lower GC sequence content on average, a feature favoring nucleosome formation (**Figure 5A**) (Deniz et al. 2011). We then asked whether these sequences were likely to form nucleosomes based on a previously reported *in vitro* reconstitution assay (Locke et al. 2013). We found that DNA sequences occupied by fragile nucleosomes in the embryo were generally less occupied *in vitro* (**Figure 5C**). Finally, we observed that sequences occupied by fragile nucleosomes were less conserved across nematodes than DNA sequences occupied by the set of all nucleosomes (**Figure 5E**).

**Figure 5.**
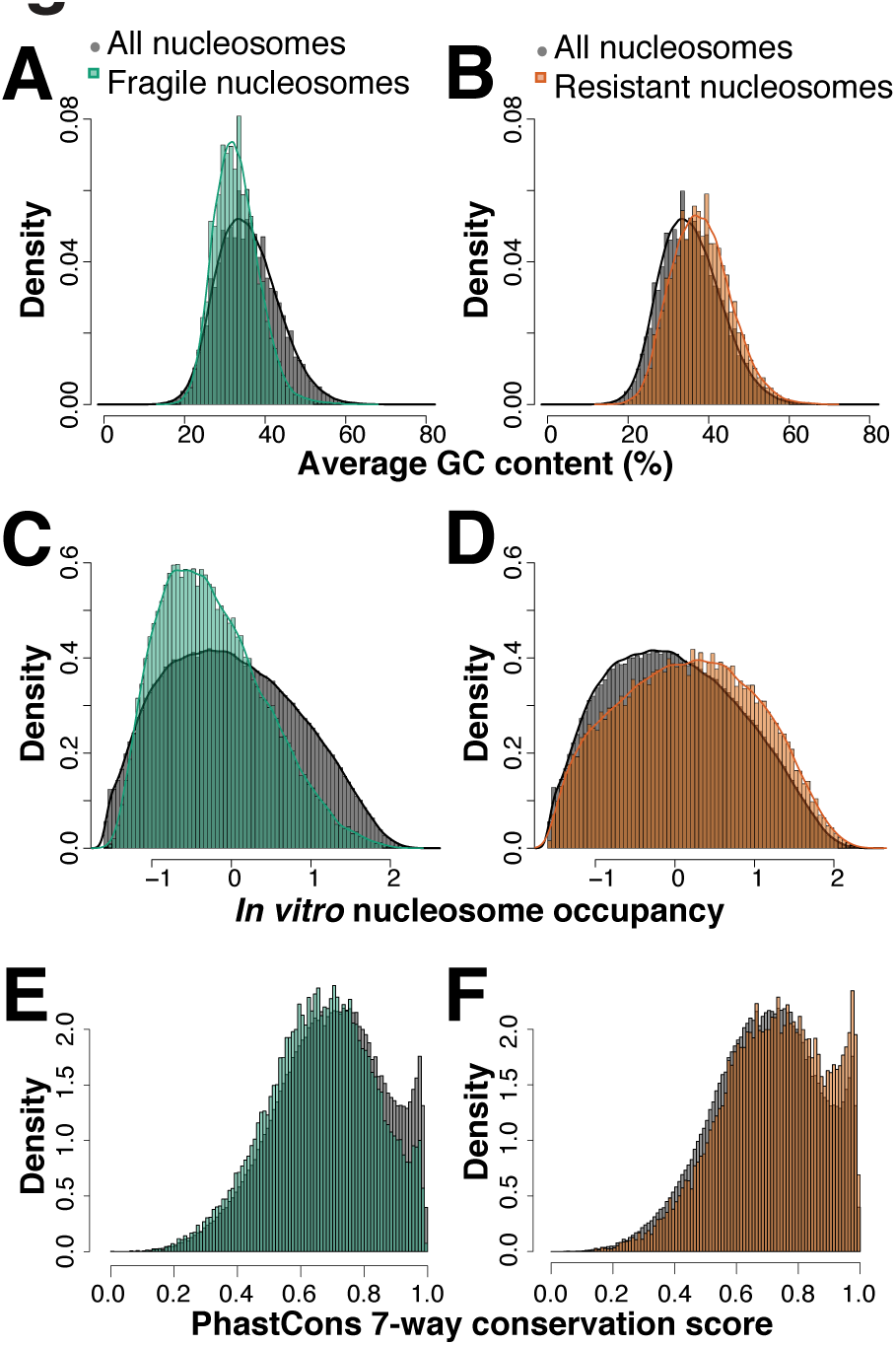
Fragile nucleosomes are found at AT-rich, nucleosome disfavoring, and poorly conserved promoters. **(A)** Histogram of average GC content at fragile (green) or all nucleosomes (grey). **(B)** Same as A for resistant (orange) or all nucleosomes (grey). **(C)** Histogram of *in vitro* nucleosome occupancy scores at fragile (green) or all nucleosomes (grey). **(D)** Histogram of *in vitro* nucleosome occupancy scores at resistant (orange) or all nucleosomes (grey). **(E)** Histogram of PhastCons 7-way conservation score at fragile nucleosomes (green) or all nucleosomes (grey). (**(F)** Same as E for resistant nucleosomes (orange) or all nucleosomes (grey).

Poly(dA:dT) tracts disrupt nucleosome formation and tend to increase transcription of downstream genes (Raveh-Sadka et al. 2012), while TATA box motifs in yeast are associated with bendable promoters sensitive to chromatin remodelers (Tirosh et al. 2007; Albert et al. 2007). In *C. elegans*, the number of T-block motifs (3 to 5 consecutive thymine nucleotides, often spaced at 10 bp periodicity) have been positively correlated with expression: genes with more than 5 T-blocks have fivefold higher expression than genes with fewer than 4 T-blocks (Grishkevich et al. 2011), presumably through a reduction in promoter nucleosome occupancy. T-blocks were not enriched at fragile or resistant nucleosomes, whereas TATA box motifs were enriched at fragile nucleosomes (**Supplemental Figure 14**).

In addition to DNA-encoded *cis* features, promoter fragility at lowly-transcribed genes may be influenced by epigenetic features associated with these nucleosomes. Through comparison with previously generated datasets, we asked whether any histone post-translational modifications, histone variants, or chromatin states were positively associated with nucleosome fragility or resistance (Ho et al. 2014; Ooi et al. 2010). Only “low signal” chromatin states and chromatin extracted with 80 mM salt (another method proposed to identify unstable nucleosomes) were associated with fragile nucleosomes (Supplemental Figures 15, 16, 17) (Ooi et al. 2010; Ho et al. 2014). Though longer linkers were weakly correlated with increased fragility levels (**Supplemental Figure 18**), the GC content of the nucleosome was the strongest predictor of overall nucleosome fragility score (**Supplemental Figure 19**). Taken together, our data indicate that fragile nucleosomes in gene promoters are correlated with high AT content and TATA box motifs. It seems that most promoters are fragile at least in part due to high AT content (see residual fragility in **Figure 2G**), but that this *cis* effect of DNA sequence becomes apparent only at sites where the observation is not confounded by TF binding.

### Fragile nucleosomes are associated with genes expressed in future or context-specific situations

To infer potential functional implications of nucleosome fragility in the developing embryo, we next asked which genes were significantly associated with fragile nucleosomes. Fragile nucleosomes were enriched at the ‐1 nucleosome and resistant nucleosomes were enriched at the +1 nucleosome (**Figure 4**). We identified two sets of genes that contain fragile and resistant nucleosomes, respectively, +/‐5 500 bp from their transcript start site. We then identified enriched GO terms in each of the gene sets (Supek et al. 2011; Al-Shahrour et al. 2004). Genes with fragile nucleosomes were enriched for GO terms related to neuronal response, immune response, and stress response genes (“sensory perception of chemical stimulus”, “defense response”, “pharynx development”, “immune system process”) (**Figure 6A**). In contrast, genes with resistant nucleosomes were enriched for embryogenesis and cell cycle related terms ("mitotic cell cycle”, “RNA processing”, “regulation of developmental process”, “organic substance transport”) (**Figure 6B**). A complementary analysis confirmed the association between high fragility scores in the promoter and genes that function in context-specific processes. Unbiased k-means clustering of promoters based on their fragility scores identified a cluster containing lowly-expressed genes with high fragility scores that are expressed in response to perception of chemical stimulus, cognition, and neurological processes (**Supplemental Figure 20**). This second class of fragile nucleosomes in the embryo is unlikely to be fragile due to the action of *trans* factors because of their anti-correlation with expression and transcription factor binding. Rather, fragile nucleosomes were associated with lowly-transcribed genes that are expressed in a future context-specific fashion during stress response or development.

**Figure 6.**
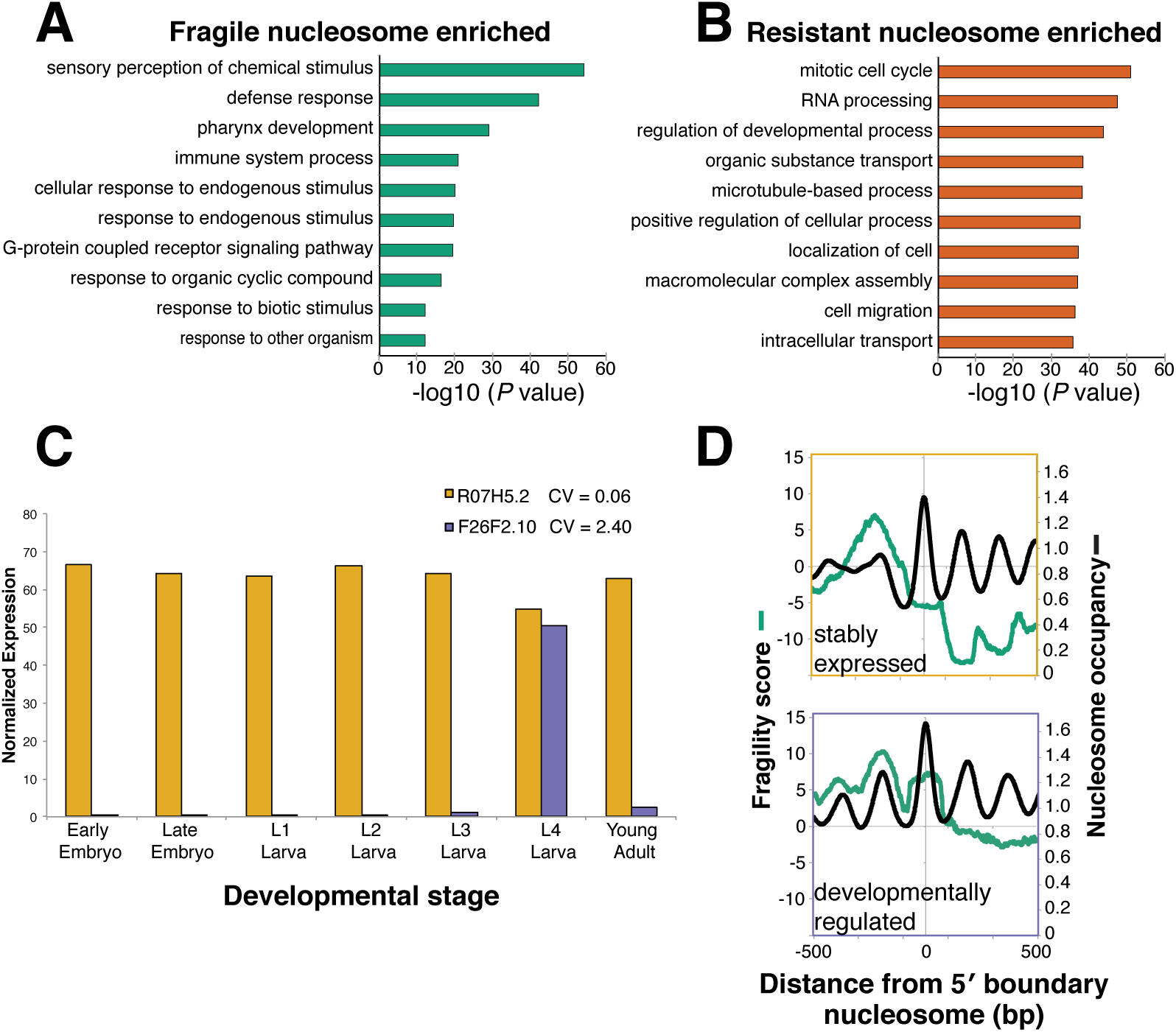
Fragile nucleosomes are enriched at genes that will be expressed in the future and in specific contexts. **(A)** Top 10 Gene Ontology biological process functional annotation terms associated with genes with fragile nucleosomes. **(B)** Top 10 gene ontology biological process functional annotation terms associated with at genes with resistant nucleosomes. **(C)** Bar plot representation of expression levels and coefficient of variation (CV) for R07H5.2 and F26F2.10. R07H5.2 has a low CV, and is an example of a stably-expressed gene; F26F2.10 has a high CV, and is an example of a developmentally regulated gene. **(D)** Average plot of fragility and nucleosome occupancy scores at 1000 stably-expressed genes (top), or developmentally regulated genes (bottom) as determined by their coefficient of variation across 7 different life stages: early embryo, late embryo, larval stages L1, L2, L3, L4, and young adult.

To confirm the association between fragile nucleosomes and future context-specific expression with an independent method, we used the publicly available modENCODE transcriptome sequencing data from seven different life stages to define a set of “developmentally regulated” genes and a set of “stably expressed” genes (**Figure 6C**) (Pérez-Lluch et al. 2015; Gerstein et al. 2014; Spencer et al. 2011). We hypothesized that if promoter nucleosome fragility is related to context-specific expression as our GO analysis suggested, then we should find higher fragility signals near developmentally regulated genes. When we plotted the average fragility scores around these genes, we indeed saw higher nucleosome occupancy and fragility signals at developmentally regulated genes as compared to the set of stably expressed genes (**Figure 6D**). While both sets of genes have fragile promoters, our data indicate that fragility is enriched at genes that tend to be expressed specifically during development, stress, or environmental stimulus response. Together, we suggest that these sequences may reflect a specialized promoter architecture that is primarily determined by high AT content, which acts to allow future disruption of nucleosome stability, and thereby the rapid induction of gene expression in a context-specific fashion (**Figure 7**).

**Figure 7.**
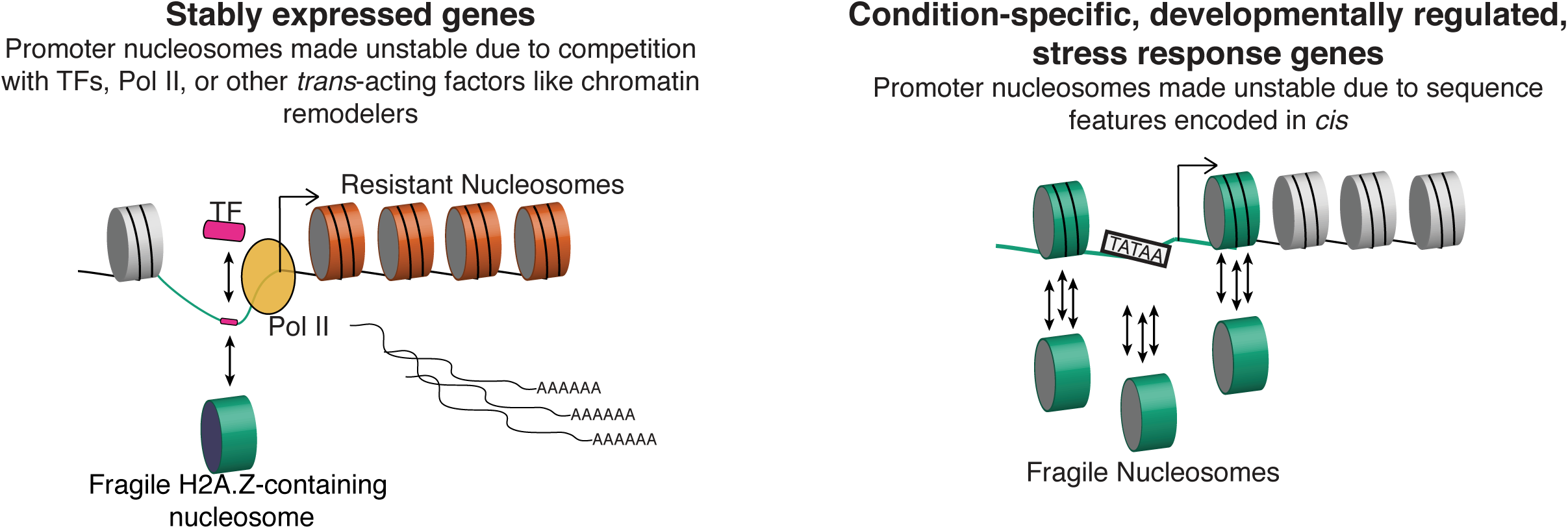
We propose a model whereby nucleosome fragility is determined by two distinct mechanisms, one that operates in *cis* at all genes, and one that operates in *trans* at a subset of genes. Left: Competition in *trans* with transcription factors and polymerase machinery destabilizes nucleosomes at the promoters of actively transcribed genes that tend to be stably expressed‥ **Right:** Condition-specific and developmentally regulated genes contain promoters with high levels of nucleosome fragility, determined primarily in *cis* by high AT content. Green line: high AT content is sequence-encoded at all promoters, but is highest at condition-specific genes. Orange cylinders: resistant nucleosomes found in the gene body of highly and stably expressed genes. Green cylinders: fragile nucleosomes compete (single arrow) with transcription factors and RNA Pol II at stably expressed genes. Fragile nucleosomes at condition-specific genes “treadmill” on the DNA (three arrows) due to destabilizing DNA elements like TATA-box motifs and high AT content.

## DISCUSSION

We performed an MNase digestion timecourse, a simple modification to the traditional MNase digestion assay, in *C. elegans* embryos. Our experiment measured which individual nucleosomes were most quickly released from their polynucleosome context after exposure to MNase. Sensitivity to MNase digestion, and thereby fragility or resistance as defined in this study, could be determined by a number of factors. These include (1) a DNA sequence in the linker region that is preferentially cut by MNase; (2) longer linker regions; (3) nucleosome instability caused by low DNA-histone affinity; or (4) instability caused by competition with transcription factors. As such, nucleosome fragility or resistance is likely to be a reasonable proxy for the underlying stability of the nucleosome. Two lines of evidence suggest that nucleosome fragility reflects nucleosome instability. First, we found that nucleosomes can be made fragile by competition with transcription factors and RNA polymerase II. Second, we observed that nucleosomes can be made unstable in *cis* by being wrapped around nucleosome-disfavoring DNA sequences with high AT content and TATA-box motifs. All of these factors have been shown to cause nucleosome instability in previous studies (Widom 2002; Ozonov and van Nimwegen 2013).

We performed our experiments in nuclei derived from whole embryos, which reflect a mixture of cell types and creates challenges for data interpretation. However, our core conclusions stand regardless. First, the DNA sequence underlying the data is the same across all cell types and therefore our conclusions regarding the *cis* contribution to nucleosome fragility are derived from and shared among all cells in the embryo. Second, previous precisely-timed developmental RNA-seq experiments show that certain genes are stably expressed in every cell type of the developing embryo, while other gene classes are not expressed at all during the time that the embryos used in our sample were collected, but instead are poised for expression later in development (Spencer et al. 2011; Gerstein et al. 2014). Therefore, for specific genes, we can say definitively that they were “on” or “off” in our sample, and make general conclusions accordingly. Third, the RNA-seq data we used is also derived from mixed embryos, and therefore quantitatively matches our fragility and resistance data. The same applies to the modENCODE chromatin data; the embryos used in this study were staged specifically to match the embryos used in the modENCODE studies. While future studies of pure populations of cells may reveal additional relationships between nucleosome properties and gene expression, the conclusions we make here with whole embryos are valid and are compatible with previously generated TF and histone ChIP-seq datasets.

### MNase resistant nucleosomes are correlated with expression in the embryo

We found a class of nucleosomes that required relatively long durations of MNase digestion to be removed from chromatin. Traditional expectations might be that unstable nucleosomes would be found in the body of transcribed genes, and stable nucleosomes in silent, heterochromatic genomic regions. However, recent reports illustrate that nucleosomes in the gene body of transcribed genes are consistently well-positioned and highly occupied due to a number of factors, including the activity of RNA Polymerase II and the histone chaperone FACT (FAcilitates Chromatin Transcription) (Bai and Morozov 2010; Jiang and Pugh 2009). Indeed, we found stable nucleosomes enriched in the gene body of actively transcribed housekeeping genes. During transcription, Pol II disrupts an H2A-H2B dimer that remains bound by FACT and is rapidly reassembled in the wake of Pol II (Formosa 2012). Histone modifications such as H3K36me3, which we found positively correlated with stable nucleosomes, are also thought to contribute to nucleosome stability and maintenance of transcription fidelity (Lieb and Clarke 2005; Lickwar et al. 2009). Together, our measurements are in agreement with an emerging picture of highly regulated nucleosome stability throughout the genome, which is likely critical for regulation of DNA templated events like transcription, splicing, and DNA replication (Bintu et al. 2011; Kwak et al. 2013; Tilgner et al. 2009; Chen et al. 2010; Eaton et al. 2010).

### MNase sensitive fragile nucleosomes are 5′ enriched and anti-correlated with expression

Differential MNase digestion and salt fractionation have been previously used to probe nucleosome-DNA stability. Results from yeast (Xi et al. 2011; Weiner et al. 2010; Kubik et al. 2015), plants (Vera et al. 2014), mouse (Lombraña et al. 2013; Deng et al. 2015), worm, (Ooi et al. 2010) and fly (Chereji et al. 2015; Henikoff et al. 2009) have identified highly labile nucleosomes in 5′ and 3′ “nucleosome free” regions. In yeast, Xi et al. observed that fragile nucleosomes were associated with H2A.Z containing promoter nucleosomes, believed to be involved in stress response (Zhang et al. 2005; Li et al. 2005). In vertebrates, nucleosomes containing both H3.3 and H2A.Z histone variants are unstable (Jin and Felsenfeld 2007; Jin et al. 2009). We did not observe a correlation between H2A.Z incorporation and nucleosome fragility in *C. elegans* as measured by our assay. Rather, H2A.Z is distribution is strongly biased towards active genes (Whittle et al. 2008; Liu et al. 2011). This distinction could be due to a divergence in H2A.Z properties between yeast and *C. elegans* (Zlatanova and Thakar 2008). It is also possible that the distribution of H2A.Z-containing nucleosomes used for this study (measured in Ho et al.) was comprised of the particularly stable, homotypic type of H2A.Z nucleosomes (Ishibashi et al. 2009).

Overall, we observed an anti-correlation between promoter nucleosome fragility and gene expression. Previous reports disagree about the relationship between nucleosome fragility and expression. Studies in yeast, *C. elegans, Drosophila*, and Maize have used salt profiling and different MNase concentrations to identify a positive correlation between promoter fragility and expression (Kubik et al. 2015; Ooi et al. 2010; Henikoff et al. 2009; Vera et al. 2014). However, Xi et al used the same approach and observed the opposite effect. One possible explanation for the discrepancy is that very light MNase digestion or salt profiling techniques may recover fragments from nucleosome-depleted loci, like the nucleosome-free regions in the promoters of active genes. If those regions are excluded, for example by slightly longer digestion times, higher enzyme concentrations, or by filtering out sub-nucleosomal fragments, truly unstable nucleosomes may become more apparent at the promoters of stress and context-responsive genes. We also point out that the relation between fragility and other methods used to measure nucleosome properties, such as profiling of salt-sensitive nucleosomes in *Drosophila* and *C. elegans*, remains unclear (Ooi et al. 2010; Henikoff et al. 2009). Both differential MNase digestion experiments and salt profiling experiments endeavor to measure some aspect of nucleosome stability but a rigorous study is needed to compare results from the two methods.

A previous study (Xi et al. 2011) identified nucleosome fragility at nearly one-third of all promoters of protein-coding genes, and enriched at the promoters of genes involved in stress response. When we assessed the types of functional annotations that were enriched at promoters with fragile nucleosomes in *C. elegans* embryos, we identified GO terms related to context-specific expression: sensory perception of chemical stimulus, defense response, immune system process. Based on our findings in conjunction of those with Xi et al., we propose that nucleosome fragility may serve to poise genes for rapid activation in response to developmental or external stimuli. This is consistent with previous work investigating the transcriptional activation of mammalian primary response genes, where unstable nucleosomes are used to achieve rapid induction independent of chromatin remodeling complexes (Ramirez-Carrozzi et al. 2009).

We found high fragility scores at the ‐2, ‐1, and +1 nucleosomes of developmentally regulated genes in comparison to stably expressed housekeeping genes. Previous work found that the promoters of developmentally regulated genes lack the histone post-translational modifications associated with active genes, like H3K4me3 (Pérez-Lluch et al. 2015). Similarly, we were unable to find an association between fragile nucleosomes and any histone post-translational modifications examined by the modENCODE group. Given the increased fragility of these nucleosomes, it is possible that (1) these nucleosomes at developmentally regulated genes were lost from standard chromatin preparation protocols and are thus underrepresented in the histone ChIP, or (2) developmentally regulated genes use promoter nucleosome fragility as a mechanism for gene regulation.

Our results are reminiscent of previous reports from yeast, which propose that promoter structures can generally be classified as containing depleted proximal nucleosomes (DPN) or occupied proximal nucleosomes (OPN) (Tirosh and Barkai 2008). In yeast, DPN genes have low transcriptional plasticity (defined as the capacity to modulate transcription levels upon changing conditions), well positioned nucleosomes, and are enriched for TF binding sites and H2A.Z. In contrast, OPN genes have high transcriptional plasticity, higher evolutionary divergence, higher nucleosome turnover, and were sensitive to chromatin regulation. The yeast DPN genes may correspond to the set of stably-expressed genes we defined in *C. elegans*, which that have depleted proximal nucleosomes (**Figure 6D** top). The yeast OPN genes may correspond to the set of developmentally regulated genes we defined in *C. elegans*, which have high promoter fragility and highly occupied proximal nucleosomes (**Figure 6D**, bottom). To our knowledge, OPN and DPN-type promoters have not been described or defined in *C. elegans*. Our results are consistent with a model in which nucleosome instability is encoded at the promoters of DPN-type genes, potentiating the high transcriptional plasticity observed at these sites. The presence of these promoter structures in yeast, human, and now *C. elegans* suggests a well-conserved strategy that uses nucleosome architecture to regulate the dynamics of gene expression.

## MATERIALS AND METHODS

### Worm strains and growth in liquid culture

Wild type *N2* worms were obtained from the *Caenorhabditis* Genome Center and maintained at 20° C in liquid culture as previously described (Whittle et al. 2008; Ercan et al. 2011). Mixed-stage embryos were isolated from gravid adults by bleach hypochlorite treatment and fixed with 2% formaldehyde for 30 minutes at room temperature.

### MNase Digestion Time Course

MNase digestion was performed as previously described (Ercan et al. 2011), with slight alterations. Micrococcal nuclease (Worthington LS004798) was resuspended in water at 50 U/uL and frozen in individual aliquots at ‐80°C. To control for variability in enzyme activity, individual aliquots were removed from the freezer and thawed on ice, and never reused. Mixed-stage embryos were incubated with chitinase (Sigma Cat #C6137), washed, and dounced in dounce buffer (0.35 M sucrose, 15 mM HEPES-KOH pH 7.5, 0.5 mM EGTA, 5 mM MgCl_2_, 10 mM KCl, 0.1 mM EDTA, 1 mM DTT, 0.5% TritonX-100, 0.25% NP-40) to extract nuclei. Nuclei were pelleted and washed with MNase digestion buffer (110 mM NaCl, 40 mM KCl, 2 mM MgCl_2_, 1 mM CaCl_2_, 50 mM HEPES-KOH pH 7.5). MNase was added, and at each timepoint (0, 2, 4, 8, 15, or 30 minutes after enzyme addition) a fraction of the reaction was removed, quenched with EDTA, and stored on ice. Samples were treated with Proteinase K for 2 hours at 55°C, then incubated overnight at 65°C to reverse crosslinks. DNA was isolated from RNA and proteins using phenol:chloroform extraction and RNase A treatment for 1 hour at 37°C. Mononucleosome-sized fragments (100 to 200 bp) were extracted from a 2% agarose gel and purified using a Qiagen gel extraction kit.

### Heat shock

Mixed-stage embryos were isolated as described and split into two pools. One pool was incubated at 34°C for 20 minutes with intermittent brief mixing, while the other pool nutated at room temperature. After 20 minutes, an aliquot from each pool was saved for RNA-seq, while the remaining embryos were fixed for 30 minutes in 2% formaldehyde at room temperature.

### RNA isolation

Embryos were dropped into TRIzol (Life Technologies) and flash frozen in liquid nitrogen after incubation for 20 minutes at room temperature or 34°C heat shock. Embryos were homogenized by thawing at 37°C and refreezing in liquid nitrogen 3x. Total RNA was isolated using a TRIzol/chloroform extraction followed by RNeasy Mini (Qiagen) preparation with On Column DNaseI Digestion (Qiagen).

### Illumina Library Preparation

Individual libraries were prepared with unique barcodes for each timepoint from the timecourse. MNaseTC libraries were prepared from 100 ng of gel-extracted DNA using the Illumina TruSeq DNA library preparation kit v2 (FC-121-2001) according to manufacturer instructions. Ampure beads were used for the final purification in lieu of gel purification. RNA-seq libraries were prepared from 2 ug of total RNA using the Illumina TruSeq RNA library preparation kit v2 (RS-122-2001) according to manufacturer instructions. Individual samples were barcoded with unique 6bp index sequences contained within the sequencing adapters. Individual libraries were then pooled at equimolar ratios for paired-end multiplex sequencing.

### Illumina Sequencing and Post-Processing

Paired-end sequencing was performed by the Princeton University Sequencing Core Facility according to Illumina protocols. Paired end reads were mapped to the UCSC Oct. 2010 (WS220/ce10) genome release using Bowtie (v1.1.2) with stringent multimapping parameters: bowtie ‐q ‐X 2000 ‐‐fr ‐p 1 ‐S ‐n 2 ‐e 70 ‐l 28 ‐‐pairtries 100 ‐‐maxbts 125 ‐k 1 ‐m 1 ‐‐un /Unmapped_Reads.fastq ‐‐phred33-quals /ce10 ‐1 /read1.fastq ‐2 /read2.fastq

### Nucleosome analysis

Reads with insert sizes between 100 and 250 bp were kept for downstream analysis.

Replicates were first processed individually, then pooled after confirming a high degree of correlation between replicates. Nucleosome analysis was performed as described previously (Kaplan et al. 2010; Gossett and Lieb 2012).

#### Coverage

Nucleosome coverage was calculated by extending the filtered mapped reads to their fragment length and measuring the sum of reads covering each bp. To normalize for variation between samples, nucleosome coverage was scaled by 1/(mean coverage), yielding a mean nucleosome coverage of 1.0.

#### Dyads

Dyads are approximated as the center of a paired-end fragment. The number of dyads at each base pair was scaled by 1/(mean dyad density), then Gaussian smoothed with a standard deviation of 20 bp.

#### Nucleosome calls

Nucleosome positions were identified from dyad density maps using a previously reported greedy algorithm (Albert et al. 2007; Gossett and Lieb 2012). Using the local maxima of the dyad density as the nucleosome center *p*, the size of the nucleosome (the nucleosome-protected region) was determined by measuring the average length of all reads that covered the nucleosome center. The standard deviation of the nucleosome center (the nucleosome “fuzziness”) was calculated for each called nucleosome as the standard deviation of dyads around the mean. Nucleosome occupancy was defined as the number of dyads that fell within 50 bp of the nucleosome center.

#### Boundary nucleosomes

Using these called nucleosome positions 5′ and 3′ boundary nucleosomes were identified for the 20,578 RefSeq annotated genes. 5′ +1 nucleosomes were identified as the first nucleosome call with a dyad coordinate downstream of the 1st coding exon. Similarly, the 3′ boundary nucleosome was identified as the first nucleosome call with a dyad coordinate upstream of the TTS. Because *C. elegans* utilizes trans-splicing, the 5′ end of mature polyadenylated mRNAs, and the RefSeq annotations used in this analysis, do not reflect the exact base pair position of transcription initiation. Although recent studies have used novel methods to identify the true transcription initiation sites (Chen et al. 2013; Kruesi et al. 2013; Saito et al. 2013), the TSS annotations are only known for a subset of expressed genes in a small number of stages (Chen et al. only tested embryos, only 31.7% of genes had a TSS in at least 1 of 3 stages tested in Kruesi et al., Saito et al., only tested embryo and adult). For completeness, we chose to instead investigate the full set of known genes using their first coding exon as an alignment point.

### Nucleosome Fragility and Resistance scores

The pooled “intermediate” nucleosome profile was generated by pooling the reads from the 4, 8, and 15 minute time points from each replicate. The pooled reads were used to generate average nucleosome positions, fuzziness scores, and occupancies. To identify regions of the genome that were liberated earlier or later than average, we subtracted the occupancy of the pooled sample from either the 2 minute (2m - pool = Fragility score) or the 30 minute samples (30m - pool = Resistance score). To highlight regions significantly enriched with this signal, we considered the 10% of nucleosomes with the highest fragility or resistance scores as Fragile or Resistant Nucleosomes.

### Gene ontology analysis

Gene lists were uploaded to the FatiGO web server (babelomics.bioinfo.cipf.es) and compared against the background set of all *C. elegans* genes (Al-Shahrour et al. 2004). P-values were calculated using the Fisher’s exact test, and corrected for multiple testing using the FDR procedure of Benjamini and Hochberg (Benjamini and Hochberg 1995). Corrected p-values and GO terms were then input in to REVIGO to reduce and visualize significantly enriched GO clusters (Supek et al. 2011).

### Stable and developmentally regulated genes

Pre-normalized transcriptome sequencing data was downloaded from: https://www.encodeproject.org/comparative/transcriptome/(Gerstein et al. 2014; Spencer et al. 2011). For each gene, we calculated the coefficient of variation (CV): 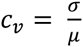 We took the 1000 genes with the highest CVs as the set of developmentally regulated genes, and the set of 1000 genes with the lowest CVs as the set of stably expressed genes.

### RNA-seq analysis

Unstranded mRNA libraries were prepared from total RNA for RNA-seq using the Illumina TruSeq RNA Library Preparation Kit v2 (RS-122-2001). RNA-seq reads were mapped to the *C. elegans* WS220 Gene Annotation Model using Tophat2 (v0.7) (Trapnell et al. 2012). The resulting alignment files were quantified using HT-Seq (v0.4.1) and the RefSeq gene annotations for WS220 (Anders et al. 2015). Total read counts per gene were normalized for differential expression using DESeq2 (v1.0.19) in R (v3.0.1) (Love et al. 2014).

### Additional datasets

A brief description of the additional publicly available datasets used in this study and their accession numbers can be found in **Supplemental Table S1**.

## DATA ACCESS

The sequencing data from this study have been submitted to the NCBI Sequence Read Archive (SRA; http://www.ncbi.nlh.nih.gov/sra) under accession number SRP072274. Additionally, processed and raw data from this study have been submitted to the NCBI Gene Expression Omnibus (GEO; http://www.ncbi.nlm.nih.gov/geo/) under accession number GSE79567.

## ACKNOWLEDGEMENTS

We thank Coleen Murphy and members of the Lieb laboratory for feedback and comments on the manuscript. This project was supported by NIH NHGRI grant U01 HG004270. TEJ was supported by NHGRI predoctoral training grant T32HG003284-10S1. Some strains were provided by the CGC, which is funded by NIH Office of Research Infrastructure Programs (P40 OD010440).

## AUTHOR CONTRIBUTIONS

Experiments were designed by TEJ and JDL. Experiments and bioinformatics analyses were performed by TEJ with guidance and feedback from JDL. Manuscript was prepared by TEJ, and edited by JDL.

